# Optimizing Communication of Emergency Response Adaptive Randomization Clinical Trials to Potential Participants

**DOI:** 10.1101/091819

**Authors:** Bredan McEvoy, David Haidar, Jason Tehranisa, William J. Meurer

## Abstract

**Introduction:** Acute clinical stroke trials are challenging to communicate to patients and families considering participation. Response adaptive randomization (RAR) is a technique that alters the proportion of trial subjects receiving active treatment, based on the outcomes of previous subjects. We aimed to determine how well interactive videos would improve understanding of a simulated acute stroke trial scenario that incorporated a design with RAR.

**Methods:** We performed a cross-sectional study of emergency department patients who were without stroke, altered mental status, or critical illness. Subjects viewed a hypothetical stroke and clinical trial scenario. They were randomized into one of four groups with either an RAR or fixed randomization clinical trial design and with either a standard consent video, or an interactive video.

**Results:** We enrolled 720 participants. In the RAR group with interactive video, 128 out of 149 (85.9%) of the subjects were able to correctly identify the allocation method, compared to the 172 out of 285 (61.6%) in the RAR group with the uninterrupted video for an absolute increase of 25.6% (95% CI 17-33%). The RAR group with interactive video had a higher odds of correct identification of allocation method (O.R. 2.767, 95% CI [1.011 – 7.570] while controlling for age, sex, ethnicity, education, self-reported understanding of protocol, stroke awareness and agreement to participate in trial.

**Conclusions:** The interactive video increased participant understanding of an RAR design in a simulated stroke scenario. Future research should focus on whether acute trial recruitment can be enhanced using similar techniques.

## Introduction

In the Emergency Department, the rapid communication of clinical research trials to potential participants can be difficult. Due to time sensitivity, preconceived notions about risks of joining trials, and a negative attitude towards research in the general public, people hesitate to participate in emergency research studies – with only about half consenting to participate.^1^ However, the introduction of random-adaptive randomization (RAR) within clinical trials has helped alleviate this problem.^2^ In trials that utilize RAR, predetermined guidelines are used to adjust the randomization ratio of participants assigned to each study group based on accumulating data in the ongoing study. This trial design allows for more patients to be assigned to a better performing treatment without compromising the scientific integrity of the study. Clinical trials typically do not improve outcomes for their participants, but implementing RAR favors patients in trials where one treatment is better than the other, and allows for an increased probability that participants can benefit from participating in clinical trials.^2^

If patients understand that they will have an increased chance of benefitting from a trial, then in the use of RAR should lead to an increase in patient participation when compared to standard fixed randomization.^3^ However, with the increased complexity of RAR, properly conveying the necessary information about the trial to potential research subjects in a clear, concise, and comprehensible manner becomes even more challenging. In 2012, we presented hypothetical stroke scenarios to 418 emergency department patients at the University of Michigan, and asked them if they would be interested in signing up for the clinical trial based on our presentation. 140 of the 208 (67.3%) in the RAR group chose to participate in this study, compared to only 114 of 210 (54.3%) of those in the standard group, with an absolute difference of 13% (95% CI: 3.7 to 22.1%). However, significantly fewer in the RAR group (62%) were able to correctly identify the method of trial allocation when compared to the standard group (85%). Thus, although participation was higher, participants were less likely to completely understand the protocol or correctly identify the allocation method if presented with the RAR model.

Our goal was to present an adjusted trial description of the hypothetical stroke scenario to emergency department patients without stroke. We hypothesized that altering the short trial description by integrating brief comprehension questions into the consent process, could improve participant understanding and therefore increase the number of individuals who can correctly identify the allocation method if presented with the RAR trial.

## Methods

### Study Design

We performed a cross-sectional study of emergency department (ED) patients at the University of Michigan between June 2 and August 1, 2014 with random allocation to two hypothetical clinical trials.

### Study Population

Patients were screened in the ED from the electronic medical record, MiChart. The study consisted of adult patients (age ≥ 18) in the University of Michigan ED who presented without symptoms of stroke or altered mental status, with stable vital signs, and who were not located in a resuscitation bay.

### Study Interventions

The subjects were introduced to the study and verbally consented. We then assessed for stroke symptom knowledge by asking the patient to identify stroke warning signs. The patients were then randomized to one of four groups, either receiving an RAR or standard hypothetical acute stroke trial, with or without four additional comprehension questions. The two groups without additional comprehension questions were exposed to the same procedure as the 418 patients from the 2012 study.^3^ These scenarios were presented to patients in video form on an iPad. Further details of the protocol are available in the supplemental material, including links to the videos.

The four comprehension questions that were added to the videos for the assigned groups addressed research procedures relevant to the consent process and trial operation. One question specifically addressed the method of randomization and allocation. The following items were available to the subjects (depending on scenario): consent form for standard trial, consent form for RAR trial, and risk pictograph for stroke thrombolysis.^4^ The clinical scenario and all other aspects of the trial were exactly the same as the 2012 study. The patient was told that “time is of the essence,” and that a decision needed to be made quickly, in order to simulate the acute trial enrollment process for stroke. If the patient had a family member or other visitor present in the room, they were asked to refrain from discussing the decision with the patient until after the scenario and data collection were completed.

After viewing the videos, the patient was given the opportunity to ask questions regarding the hypothetical research trial. The patient was then asked whether or not they agreed to participate in the hypothetical trial. The patient was also asked to identify the method of randomization and allocation used in the trial. A modified version of the ICQ-4 instrument (the fourth item– which asked if the study met expectations - was excluded because this was a hypothetical study and the patients did not actually participate) was used to assess adequacy of informed consent.^5^ Patient demographics were collected upon completion of the interview and the patient was given a handout on stroke warning signs at the end of the research procedures. No protected health information or any other specific identifiers were collected from the patients.

### Study Endpoints/Outcome

The primary outcome was the difference in proportions for correct identification of the hypothetical trial allocation method between the comprehension questions (intervention) group versus the uninterrupted video group, limited to the subjects assigned to the RAR groups.

The pre-planned secondary outcome was a difference in differences analysis. The outcome of interest was the participation in the hypothetical trial (the same primary outcome as the 2012 study).

### Statistical Analysis

For the primary analysis, the 218 subjects from the 2012 study assigned to the RAR group were included in the analysis. As a pre-planned secondary analysis, we used multivariable logistic regression to estimate the adjusted odds of correctly identifying the allocation method within the RAR group, and included the following covariates based on our a priori belief about potential confounders: age, sex, ethnicity, and education. We tested for heterogeneity between the 2012 and 2014 uninterrupted video groups using chi-square.

For secondary outcome analysis focusing on agreeing with the study, all 2012 subjects were included. Proportions and 95% confidence intervals for each of the four groups were calculated with 2014 and 2012 groups (uninterrupted video) combined. Logistic regression was conducted with the following indicator variables as covariates: RAR trial versus standard, comprehension video versus uninterrupted, and interaction term. In addition, an adjusted model was fitted to include additional covariates based on our a priori belief about potential confounders: age, sex, ethnicity, education, self-reported understanding of protocol, ability to correctly identify allocation technique, and stroke awareness. Descriptive statistics for demographics and stroke knowledge were calculated. Summary scores based in the ICQ-4 scale were calculated and comparisons were made for trial accepters comparing the standard group and the RAR trial group.

### Sample Size Calculation and Randomization

We planned to enroll approximately 300 subjects. We believed this to be feasible as 418 interviews were conducted during the previous 2012 study and our protocol was not significantly lengthened in terms of time. There was no pre-specified maximum number of subjects as this was a time-limited summer project. The correct identification proportion in the 2012 study was about 62%. We calculated that if we could achieve 150 new subjects in the enhanced video group, and 75 new subjects in the uninterrupted video group (added to the 208 from 2012), we would have 90% power at the 0.05 significance level to detect an increase in the correct identification proportion to 77%, which would be clinically meaningful. (For reference, the standard randomization or “coin-flip” group from 2012 correctly identified the randomization technique 85% of the time).

The patients were randomized in balanced, randomly permuted blocks (sizes 12 and 24) in order to maintain ongoing numerical balance between the 4 groups throughout the study. The randomization scheme was generated using the website Randomization.com http://www.randomization.com. The research assistant was only able to access the assignment of the current patient (in order to properly administer the scenario) and did not have access to results within the database during the data collection phase.

### Human Subjects Protection

The study protocol was reviewed and approved by the University of Michigan Institutional Review Board and was granted Exempt status. Verbal consent was obtained from the patient to participate in the simulation and they received a handout regarding their research participation and its voluntary nature. Since this was minimal risk research - and an informed consent form would represent the collection of personal identifiers - formal written informed consent was not obtained.

## Results

### Study Population

720 participants were enrolled in the study, 418 from the 2012 study and an additional 302 from 2014. The flow of subjects is described in the supplemental material (supplemental table 1). There was not heterogeneity between the 2012 and 2014 RAR uninterrupted video groups (p=0.898, supplemental table 2). Sex, history of stroke, hypertension, diabetes, atrial fibrillation, heart attack, education, ethnicity, and previous knowledge of stroke were comparable across groups (supplemental table 3). The standard and RAR groups with interactive video were significantly older than the groups without interactive video because in 2014 we targeted older patients to participate in the trial so that our population greater reflected the patient population at risk for stroke.

### Primary Outcome

When patients were presented with a hypothetical acute stroke study, 128/149 (85.9%) of the subjects in the RAR group with interactive video were able to correctly identify the allocation method, compared to the 172 out of 285 (61.6%) in the RAR group with the uninterrupted video for an absolute increase of 25.6% (95% CI 17-33%). When limiting the analysis of the primary outcome to only subjects enrolled in the 2014 RAR groups, there is a similar, also significant 26.2% (95% CI 14-38%) absolute increase in correct allocation method identification in the interactive video group relative to the uninterrupted video. Correct identification generally was highest in the RAR interactive video groups across demographic subgroups (supplemental table 5)

### Secondary Outcomes

When patients were asked if they would agree to participate in the clinical trial, 83.3% (95% CI 77-88%) of subjects in the RAR group with interactive video agreed, compared to only 67.4% (95% CI 62-73%) of patients in the uninterrupted RAR group.

There was no significant difference in comprehension summary scores (interactive video questions answered correct out of a total of 4) when comparing standard CT with interactive video to RAR with interactive video (p-value 0.216) (supplemental table 5). There was also no significant difference in comprehension summary scores in trial acceptors between the two groups (p-value 0.42).

### Multivariable models and stratified analyses

In the multivariable logistic regression model, the RAR group with interactive video (interaction term) had a higher odds of correct identification of allocation method (O.R. 2.767, 95% CI [1.011 – 7.570]) while controlling for age, sex, ethnicity, education, self-reported understanding of protocol, stroke awareness (identify 2+ stroke risk factors) and agreement to participate in trial (supplemental table 6). The main effect of interactive video was not significant in the multivariable model, due to the high ability to identify the allocation method (coin-flip like procedure) in the standard clinical trial, with or without the interactive video. However, patients who identified as white were more likely to correctly identify the allocation method when compared to those who identified as non-white (O.R. 1.705; 95% CI 1.157 – 2.512). In stratified analyses, those who reported complete understanding were significantly more likely to correctly identify the method of allocation (79.40% versus 64.70%; OR 2.097, 95% CI 1.491 – 2.949) (supplemental table 7). In addition, we assessed the association between race/ethnicity and level of education – there was a small association noted (supplemental table 8).

## Discussion

We found that when subjects in the RAR group were presented with an interactive video, they were more likely to correctly identify the allocation method for their trial. This supports our hypothesis that the implementation of brief comprehensive questions in the consent process improved patient understanding. We also found that subjects presented with an interactive video were also more likely to agree to participate in the clinical trial suggesting that the improved understanding will help improve recruitment and increase participation in clinical trials. Our data is consistent with other studies that have shown that the implementation of the RAR method alone increases research participation when compared to the standard randomization model.^6^ However, by adjusting the RAR consent model to include interactive, comprehension testing videos, we can quickly achieve comparable understanding of the design to a much simpler, coin-flip randomization trial.

Non-white subjects were less likely to correctly identify the allocation scenario, even after adjusting for education. This could be related to the way our questions were phrased or how the video was presented. Future studies could revise the presentation of information in order to further improve patient understanding for a wider more diverse population. One method could be to implement adaptive education, in which subjects who struggle to answer more difficult questions correctly will be given more detailed descriptions to help them answer subsequent questions. Given that those who reported complete understanding had higher odds of correct allocation, and those who correctly identified allocation method were more likely to participate in clinical trials, future studies can investigate the relationship between complete understanding and agreement to participate in clinical trials. In addition to implementing interactive videos, we can potentially explore alternative methods to improve understanding in order to increase participation in future trials.

This study has some limitations. Given that the study took place in one community, a suburban academic medical center, one cannot extrapolate this information and apply it generally. Another limitation was that we presented a hypothetical stroke scenario. In an actual stroke case, patients would likely be cognitively impaired and unable to provide consent on their own.^1^ However, previous studies have indicated that willingness to participate in a hypothetical trial is a positive predictor of participation in an actual trial.^7^ This information, along with the assumption that next of kin is expected to choose what the patient wants, and that studies show that surrogates tend to make decisions that they themselves would choose, one can reasonably believe these results will most likely reflect reasonable results for proxies as well.^1^ Finally, the two arm RAR trial with a binary outcome will lose statistical efficiency but potentially improve the outcomes of subjects by assigning more to the better arm (should one actually exist). Adaptive trials incorporating RAR may have more aligned statistical and clinical benefits when they have more than 2 arms, an important example is the Established Status Epilepticus trial.^8^

In conclusion, our findings suggest that the implementation of comprehensive videos when obtaining consent will further improve understanding in an acute clinical trial using RAR. We also found a further increased rate of participation in the RAR trial when compared to the standard trial. Thus, such methods could be incorporated into ongoing or planned future clinical trials in order to improve subject participation which could lead to more efficient future clinical trials with faster and more inclusive enrollment. Previous studies have shown that research participants are uncomfortable with random chance determining their treatment, and that they would prefer to receive the new treatment.^9^ Given this information, it is in our best interest to continue exploring the RAR method in order to increase research participation. Our next step could be to possibly implement this project in multiple hospitals to further diversify our study population to improve the generalizability of this study. Decision making in a time sensitive setting such as the emergency department also plays a role in the choices participants make. Future studies could vary the amount of time given to participants to assess if this changes one's decision to agree to participate in clinical trials. The ability to balance giving one enough time to process the information while still making a timely decision is difficult but an important skill to improve upon in order to improve future studies. Improving understanding in the emergency setting will ideally lead to a more positive perception of clinical trials and lead to an increased participation in future trials.

## Data Access and Responsibility

Dr. Meurer had full access to the data and takes full responsibility for the analysis and results. The analytic dataset is available for download at this digital object identifier doi:10.7302/Z2XS5S9J

## Potential Conflicts of Interest

The authors report no conflicts of interest

## Financial Support

Funding from the University of Michigan Medical School Student Biomedical Research Program funded by the National Institutes of Health 5T35HL007690-30

